# Mechanical force induces DRP1-dependent asymmetrical mitochondrial fission for quality control

**DOI:** 10.1101/2022.10.27.513965

**Authors:** Xiaoying Liu, Linyu Xu, Yutong Song, Xinyu Li, Cheuk-Yiu Wong, Rong Chen, Jianxiong Feng, Hei-Man Chow, Shuhuai Yao, Song Gao, Xingguo Liu, Liting Duan

**Author notes:** Correspondence (L.D).

## Abstract

Mitochondria are membrane-bound organelles that perform diverse critical biological functions. They undergo constant fission and fusion, which are important for mitochondrial inheritance, functions, and quality control. While tremendous efforts have identified many factors governing mitochondria dynamics, emerging evidence indicates the involvement of various intracellular or extracellular mechanical cues. However, how mechanical stress directly modulates mitochondrial dynamics remains largely unknown. Here utilizing an optogenetic mitochondria-specific mechanostimulator to apply pulling forces to intracellular mitochondria, we find that mechanostimulation can promote mitochondrial fission, with sustained mechanostimulation triggering fission more effectively than transient one. Asymmetrical fission can occur at different sub-mitochondrial sites after force-induced mitochondrial elongation. Such force-induced fission is dependent on DRP1 and involves the wrapping of ER tubules. Moreover, mechanical force generates mitochondrial fragments without mtDNA which recruit Parkin proteins. Our results prove the mechanosensitivity and mechanoresponsiveness of mitochondria and reveal the role of mechanical cues in directly regulating mitochondrial dynamics.

## Introduction

Mitochondria are membrane-bound organelles present in almost all eukaryotic cells. They execute and coordinate many essential cellular processes, including cellular energy production, calcium buffering, reactive oxygen species generation, and biosynthesis of steroid hormone and heme. Mitochondrial dysfunction severely affects cell homeostasis and contributes to a variety of diseases ranging from cancer, neurodegeneration, cardiovascular disease, to metabolic disorders^1^. As highly dynamic organelles, mitochondria continuously undergo fusion and fission, which shape the mitochondrial network, control inter-mitochondrial interactions, and maintain mitochondrial functions^2^. Moreover, by splitting damaged mitochondrial components into daughter cargos for further repair or degradation, mitochondrial fission is employed as an important quality control pathway to safeguard the vital functions of mitochondria^3^. Enormous efforts have been dedicated to uncovering the diverse mechanisms regulating mitochondrial fission, which, however, are not completely understood to date.

Emerging evidence indicates mechanoregulation of mitochondrial functions. Mitochondria are constantly exposed to mechanical forces that are applied to cells and organisms and then transmitted to intracellular mitochondria via the crowded cytosol and cytoskeleton. On the other hand, mitochondria receive various mechanostimulation naturally generated inside the cells, such as forces engendered by molecular motors that are recruited to the mitochondrial membrane, and forces obtained by hitchhiking on other organelles^4,5^. It has been shown that mechanosignalling can regulate the biogenesis and bioenergetics of mitochondria^6^. Moreover, accumulating reports suggest the contribution of mechanical cues in promoting mitochondrial fission. Stiff extracellular matrix can induce mitochondrial fragmentation^7^. Mechanical stretching or increased pressure exerted on cells in vitro^8-10^, and mechanical ventilation or mechanical needle stabbing of cells in vivo^11,12^ have been shown to increase mitochondrial fission and fragmentation. Moreover, intracellular mitochondrial fission can be triggered by forces from a motile bacterium colliding against mitochondria, mechanical pressure created by atomic force microscopy, or constrictive force generated by uneven surfaces^13^. However, much remains obscure about whether and how mechanical forces directly regulate mitochondrial fission.

Here by utilizing an optogenetic mechanostimulator for intracellular mitochondria^14^, we investigated the connections between mechanical forces and mitochondrial fission. By optically recruiting molecular motors onto mitochondrial membranes via genetically encoded optical hetero-dimerizers, this optogenetic strategy uses blue light signals to direct the pulling forces generated by molecular motors to mitochondria. Thus, specific mechanostimulation of mitochondria in live cells can be achieved with remote control, high-throughput, and spatiotemporal precision. We found that mechanical stimulation can promote the division of mitochondria, and sustained force application can lead to more fission events than transient one. After the exertion of pulling forces and the subsequent mitochondrial elongation, asymmetric fission can be triggered at distinct sub-mitochondrial sites. We revealed that such force-elicited fission is dependent on DRP1 and involves the wrapping of ER tubules on the fission site. Asymmetric segregation of mitochondrial DNA (mtDNA) into daughter mitochondria can be induced, generating mitochondrial fragments which contain no mtDNA and later recruit Parkin proteins. Our results provide both direct proof for and mechanistic insights on the mechanostimulation-triggered mitochondrial fission, highlighting the importance of mechanical cues in modulating mitochondrial functions and strengthening the notion that the mitochondrion, as a mechanosensor, can actively contribute to cellular mechanosensing and mechanotransduction.

## Results and Discussion

### Mechanical stimulation promotes mitochondrial fission

To probe the relationship between mechanostimulation and mitochondrial fission, we used our recently developed optogenetic strategy to exert pulling forces specifically towards intracellular mitochondria^14^. In our design, the optical dimerizer iLID (improved light-induced dimer) and the micro variant of SspB were utilized which bind together within seconds after blue light exposure (Figure 1A). iLID was fused with tdTomato and the transmembrane domain of Omp25 (outer membrane protein 25, a.a 111-145) which anchors to the outer mitochondrial membrane (OMM)^15^ to target and fluorescently label mitochondria. A truncated version of KIF5A which is unable to bind to cargos was fused with SspB^16^. Upon blue light illumination, iLID/SspB interaction leads to the recruitment of kinesins to OMM, thus imposing pulling forces on mitochondria.

**Figure 1.**
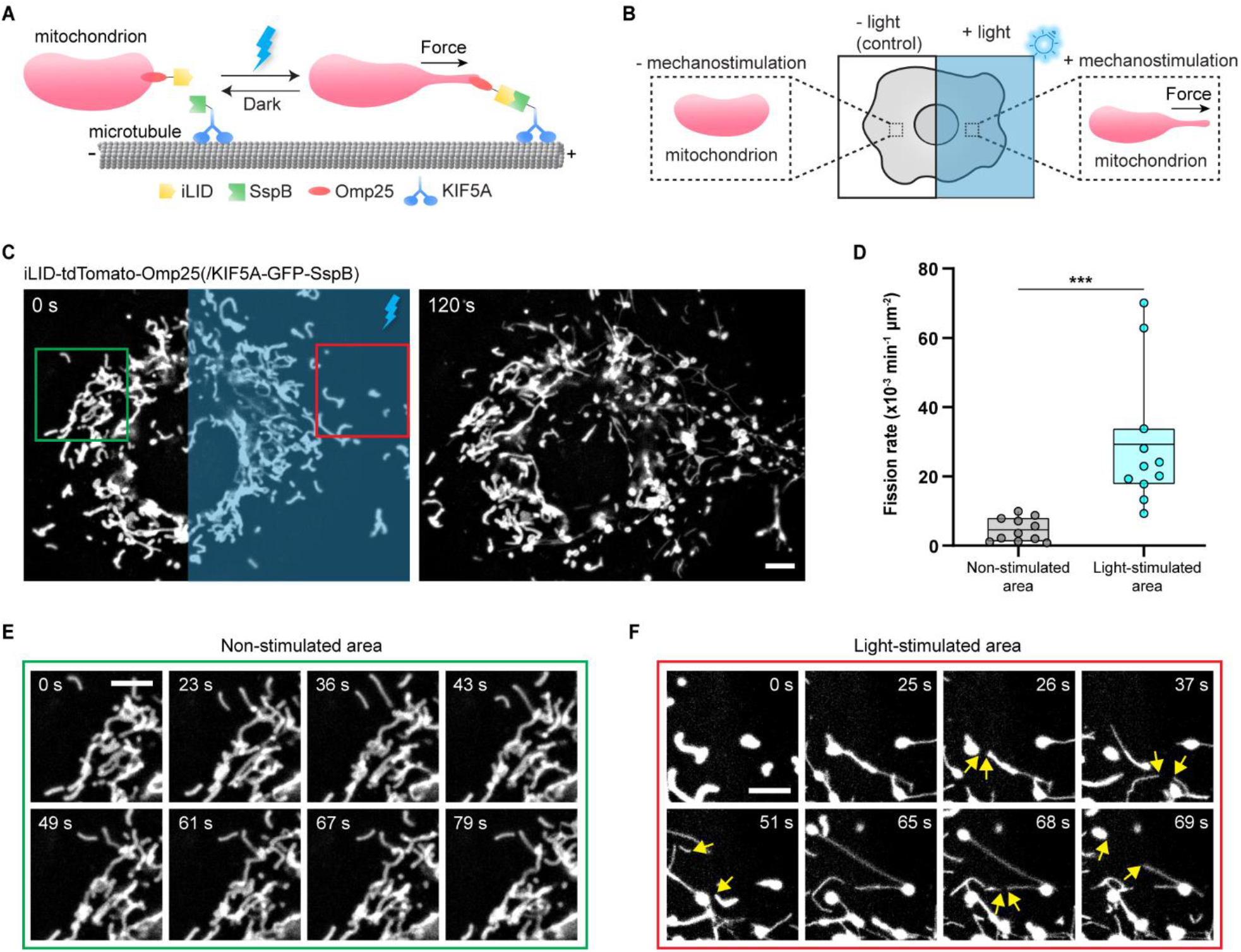
Mechanical forces by optogenetic mechanostimulator promotes mitochondrial fission. (A) Schematic representation of light-gated mechanostimulation imposed on mitochondria. iLID is fused to Omp25 to target mitochondrial membrane, while SspB is fused to the kinesin motor protein encoded by KIF5A. Blue light-gated iLID/SspB dimerization induces the recruitment of kinesins to mitochondria, thus exerting pulling force on mitochondria. (B) Schematic representation of experimental design. Blue light is delivered to one side of the cell to apply mitochondria-specific mechanostimulation, and the other side is used as a non-stimulated control. (C) Fluorescence images of tdTomato-tagged mitochondria in the COS-7 cell expressing iLID-tdTomato-Omp25 and KIF5A-GFP-SspB. The blue rectangle-marked area received intermittent blue light stimulation for 120 s. (D) Quantification of mitochondrial fission rates in the blue light-illuminated and non-illuminated areas of the same cell. N = 11 cells from 5 independent experiments. Line, mean; bounds of box, 25th and 75th percentiles; whiskers, minimum to maximum value; circles, individual values. *\*\*\*p* < 0.001 using two-tailed paired Student’s t-test. (E) Zoomed-in images of the region marked by the green box in the non-stimulated area in (C). (F) Zoomed-in images of the region marked by the red box in the blue light-stimulated area in (C). Fission events are shown by yellow arrows. Scale bars, 5 μm.

Our optical method reconstitutes one type of intrinsic mechanostimulation toward mitochondria, during which molecular motors naturally bind to mitochondria and pull the protrusion of mitochondrial tubules^4^. Moreover, we observed that some mitochondria underwent fission soon after the natural formation of thin mitochondrial tubules (Figure S1A), in concordance with the previous report^17^. Therefore, we hypothesized that mitochondrial fission can be triggered by the pulling force controlled by our optogenetic method. COS-7 cells were transfected with iLID-tdTomato-Omp25 and KIF5A-GFP-SspB. To test our hypothesis, we took advantage of the spatial precision offered by the optogenetic switch and optically exerted forces on mitochondria residing within one side of the cell, while mitochondria on the other non-illuminated side remained unperturbed as a control (Figure 1B). As shown in Figure 1C, the morphology of mitochondria within the non-illuminated area remained largely unaltered. Contrastingly, in the blue light-illuminated region marked by a blue rectangle, the mitochondrial network became more fragmented as a result of active mitochondrial fission. Close examination of the non-stimulated area marked by the green rectangle showed no noticeable deformation of mitochondria or fission (Figure 1E). However, in the stimulated area indicated by the red rectangle, thin and long mitochondrial tubules were pulled out which later underwent fission as indicated by the yellow arrows (Figure 1F). Quantification of mitochondrial fission rates in the stimulated and unstimulated areas from 11 cells in 5 independent experiments confirmed the significantly increased level of mitochondrial fission in the light-illuminated areas compared with the non-illuminated regions (Figure 1D). Therefore, our results prove that mechanical force can promote mitochondrial fission.

### Sustained force promotes mitochondrial fission more efficiently than transient force

Next, we probed how different levels of force exertion will affect mitochondrial fission differently. In our and others’ observations, some naturally extending mitochondrial tubules were retrieved back towards the mitochondria body instead of being divided (Figure S1B)^17^. We speculated that the distinct fates of mitochondria following the mechanostimulation might emanate from different temporal profiles of the force application. To probe this hypothesis, we leveraged the temporal precision of our optogenetic methods to apply transient or sustained pulling forces to intracellular mitochondria. Such precise temporal controllability relies on the reversible light-gated association between iLID and SspB so that blue light retrieval prompts the dissociation of engineered motors from mitochondria and thus abrogates the light-mediated force application. Therefore, a longer duration of blue light illumination empowers more sustained mitochondria-targeted mechanostimulation. In cells expressing iLID-tdTomato-Omp25 and KIF5A-GFP-SspB, to impose a transient mechanical force, one 200 ms pulse of blue light was delivered, while intermittent blue light exposure for 5 min was adopted for a more sustained mechanostimulation (Figure 2A). To measure the effects of light-gated mechanostimulation, mitochondria dynamics were monitored 5 min before and after the onset of blue light illumination.

**Figure 2.**
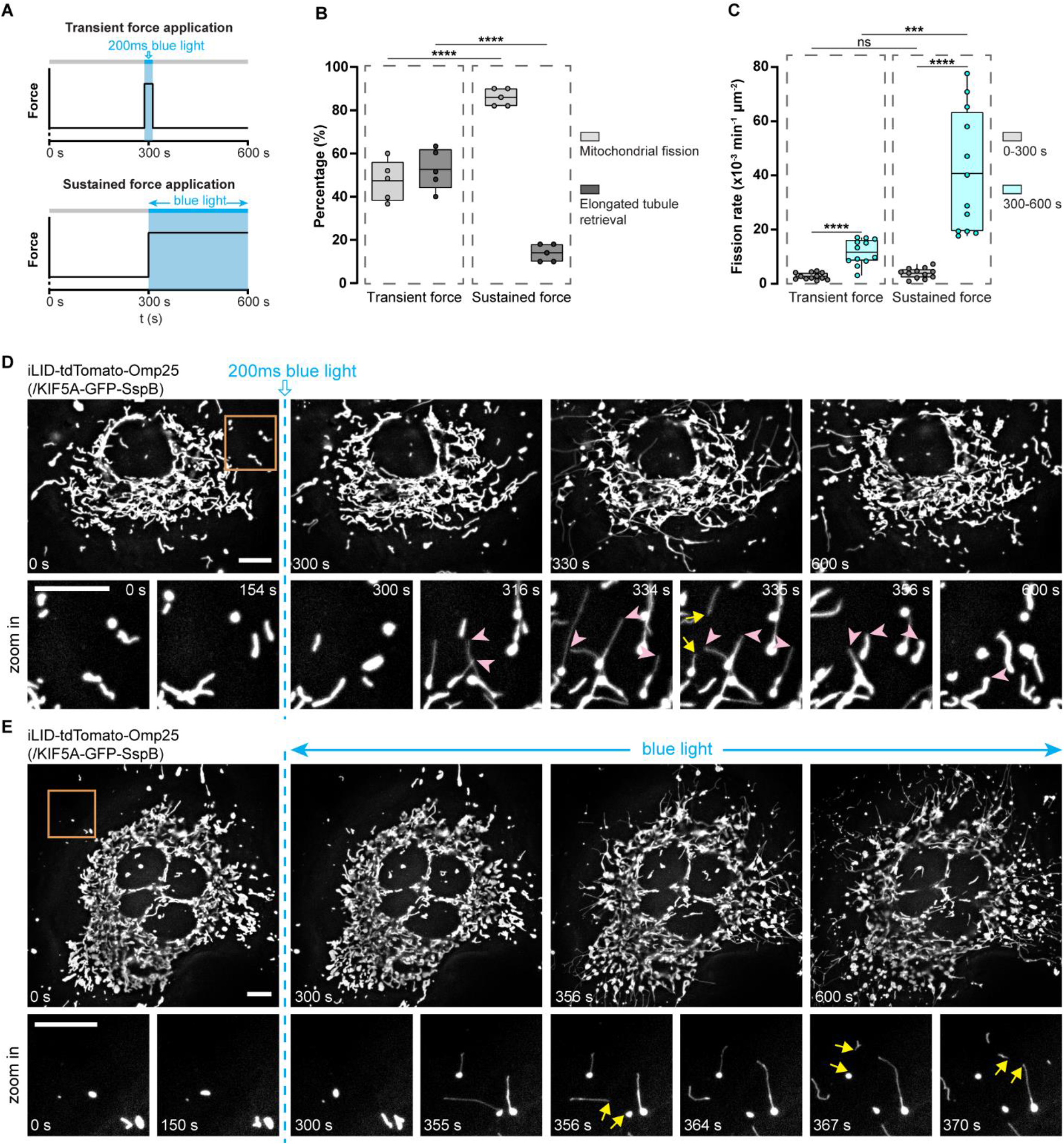
Sustained force promotes mitochondrial fission more efficiently than transient force. COS-7 cells were transfected with iLID-tdTomato-Omp25 and KIF5A-GFP-SspB. (A) Illustration for transient or sustained force application by blue light. One 200 ms pulse of blue light is delivered to impose a transient mechanical stimulation, while intermittent blue light exposure for 5 min is adopted for a more sustained mechanostimulation. (B) The percentage of protruding mitochondrial tubules undergoing fission or retrieval after blue light-mediated transient or sustained mechanostimulation. N ≥ 250 force-induced mitochondrial tubulation from 5 cells per group. Line, mean; bounds of box, 25th and 75th percentiles; whiskers, minimum to maximum value; circles, values in individual cells. \*\*\*\**p* < 0.0001 using two-tailed unpaired Student’s t-test. (C) Quantification of fission rates in the same cell before (0-300 s) and after (300-600 s) the onset of light stimulation. N = 12 cells from 4 independent experiments per group. Line, mean; bounds of box, 25th and 75th percentiles; whiskers, minimum to maximum value; circles, individual values. ns, not significant (*p* > 0.05), \*\*\**p* < 0.001, \*\*\*\**p* < 0.001 using two-tailed paired Student’s t-test for the same condition, and two-tailed unpaired Student’s t-test for different conditions. (D) Fluorescence images of mitochondria before and after transient light stimulation. Zoomed-in images of the orange box-marked area showed many mitochondrial tubulations and then retrieval of the tubules (indicated by pink arrowheads), and one fission event (marked by yellow arrows). (E) Fluorescence images of mitochondria before and after sustained light stimulation. Zoomed-in images of the orange box-marked area showed elongation of the three mitochondria and their subsequent fission (marked by yellow arrows). Scale bars, 10 μm.

As shown in Figure 2D, a transient pulling force induced mitochondrial elongation and some mitochondrial fission events, which did not induce significant fragmentation of the entire mitochondrial network. The zoomed-in examination of the area highlighted by the yellow rectangle showed that after the force-triggered initiation of mitochondrial tubulation, several tubules were withdrawn to the mitochondrial body (indicated by the pink arrowheads). Only one underwent fission as marked by the yellow arrows. On the opposite, for cells exposed to 5 min blue light illumination, the sustained mechanostimulation induced dramatic mitochondria deformation and frequent fission, which resulted in noticeable mitochondrial fragmentation (Figure 2E). As shown in the zoomed-in area, all the three mitochondria remained mostly unchanged before light stimulation. Contrastingly, blue light exposure drove the protrusion of mitochondrial tubules and the subsequent fission of all three mitochondria. Compared to transient force, sustained force application leads to a higher proportion of tubule-protruding mitochondria undergoing fission rather than tubule retrieval (Figure 2B). We calculated the mitochondrial fission rates in the same cell before (t = 0-300 s) and after (t = 300-600 s) the start of blue light stimulation. The quantification indicates that although both transient and sustained mechanostimulation increased the rates of mitochondrial fission, sustained force application is much more efficient in promoting mitochondrial fission (Figure 2C).

We further explored the mechanostimulation-induced fission with other mitochondria targeting sequences, optical hetero-dimerizers, and motors as well as in other types of cells (Figure S2). We found that the utilization of another mitochondria targeting sequence, mitochondrial Rho GTPase 1 (Miro1), other optical hetero-dimerizing pairs (CRY2/CIBN and LOVpep/ePDZ), and other motor proteins (kinesin 3 motor KIF1A, a.a 1-383), can also effectively elicit the force-induced mitochondrial fission. In addition, we have demonstrated that mechanostimulation-induced robust mitochondrial fission can be observed in U2OS cells. Thus, our results show that the force-triggered mitochondrial fission is independent of expressing any components in our optogenetic system and can be achieved in different types of cells.

### Mechanical force can induce asymmetrical mitochondrial fission at different sites for different times and cause the ring closure of the donut shape

During the force-induced mechanostimulation, we observed different phenomena of mitochondrial fission. First, fission can occur at different sub-mitochondrial sites. Upon blue light-mediated mechanostimulation, usually one thin tubule was extruded from the mitochondrial body, forming three distinct regions, including the main body, the stretching tubule, and the junction connecting the body and the tubule (Figure 3A). We found that the force-induced disconnection always occurred along the tubules, either at the body-tubule junction or in the middle of the tubule (Figure 3C). The quantification of the ratio of fission events at different locations shows that fission occurring on tubules accounted for 47.4% of all the fission events, while 52.6% took place at the junction site (Figure 3B). Such differential fission sites were also observed following the naturally occuring mitochondrial tubulations (Figure S1A). None of the fission events took place in the mitochondrial body, showcasing the strong preference for mitochondrial fission in the narrow tubules rather than round bodies.

**Figure 3.**
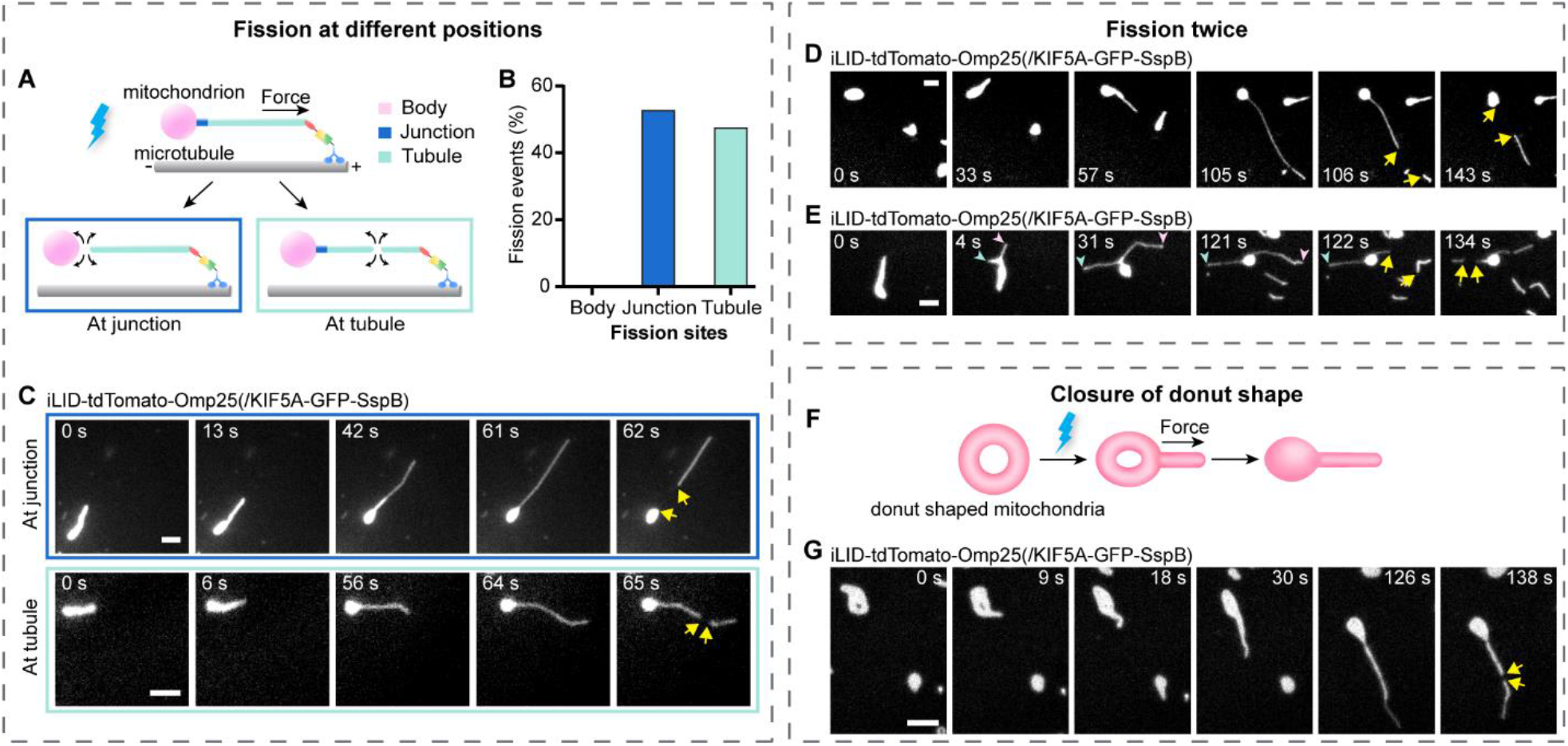
Mechanical force can induce asymmetrical mitochondrial fission at different sites for different times and cause the ring closure of the donut shape. COS-7 cells were transfected with iLID-tdTomato-Omp25 and KIF5A-GFP-SspB. (A-C) Force can induce mitochondrial fission at different sub-mitochondrial sites. (A) Illustration of force-induced fission at the body-tubule junction or in the middle of the tubule. (B) Quantification of the percentage of fission events occurring at different sites. N = 114 fission events from 11 cells. (C) Fluorescence images of mitochondria undergoing fission (indicated by yellow arrows) either at the junction or at the tubule. (D-E) Fluorescence images of mitochondria undergoing force-induced fission for twice. (D) Fission occurred twice at the same extending tubule. (E) Fission occurred at either of the two tubules protruding from the same mitochondria. (F-G) Force induced the ring closure of donut-shaped mitochondria. (F) Illustration of the force-induced ring closure. (G) Fluorescence images of donut-shaped mitochondria undergoing ring closure before force-induced fission. Scale bars, 2 μm.

Moreover, we found that one mitochondrion can experience two fission events shortly after blue light stimulation (Figure 3D, S3A). In addition, upon the formation of two extending mitochondrial tubules from the same mitochondrial body, each tubule could undergo one fission event (Figure 3E, S3B). Interestingly, when the donut-shaped mitochondria were subjected to light-mediated mechanostimulation, the ring structure was closed first and then followed by fission (Figure 3F, G, and S3C). The donut-shaped mitochondria were remodeled from tubular mitochondria under hypoxia-reoxygenation stress^18^ or increased mtROS production^19^. Upon examining the natural dynamics of donut-shaped mitochondria in cells, we observed that the donut structure could be closed shortly after mitochondrial tubulation (Figure S1C, D). Our results indicate that mechanical force can be an inherent player actively modulating the intricate mitochondrial morphology and topology.

### Force-induced fission depends on DRP1

Here we investigated whether dynamin-related protein 1 (DRP1) is involved in force-induced mitochondrial fission. As the master regulator for mitochondrial fission, DRP1 is recruited to the mitochondrial surface, forming an oligomeric ring to drive fission^20^. To explore the participation of DRP1, we transfected COS-7 cells with GFP-fused DRP1 (GFP-DRP1) as well as the optogenetic constructs (iLID-tdTomato-Omp25 and KIF5A-SspB) (Figure 4A). After blue light illumination, we found that shortly after DRP1 foci were formed on the mitochondrial tubule or junction (Figure 4B, C, and S4A, B), mitochondria were divided at these sites and then the DRP1 foci soon disappeared. Quantification of GFP-DRP1 intensity on mitochondria confirmed the enrichment of DRP1 on fission sites compared to non-fission sites (Figure 4D), suggesting that DRP1 plays a role in mechanically induced fission.

**Figure 4.**
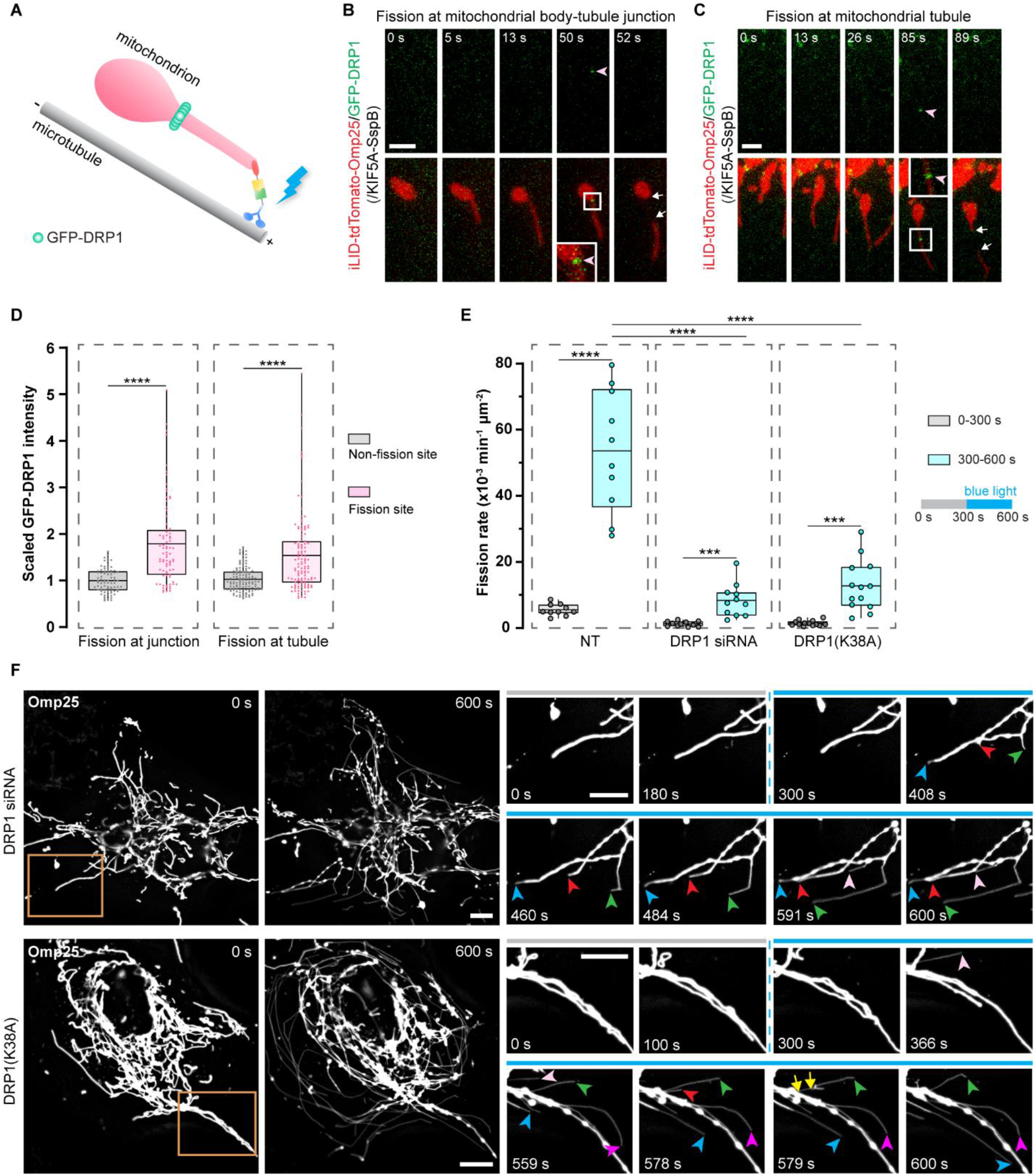
Force-induced fission depends on DRP1. (A) Illustration scheme showing DRP1 assembly at force-induced mitochondrial fission sites. DRP1 was fused with GFP to mark its distribution. (B-D) The formation of DPR1 puncta at the force-induced fission sites. COS-7 cells were transfected with iLID-tdTomato-Omp25, KIF5A-SspB, and GFP-DRP1. DRP1 puncta (pointed by the pink arrowhead) was formed at the fission site at the junction (B) or at the tubule (C). (D) Quantification of DRP1 intensity at the non-fission and fission sites on the same mitochondrion. N ≥ 80 fission events per group. Statistical analysis, Wilcoxon Matched-Pairs signed ranks test for the fission and non-fission sites on the same mitochondrion. (E-F) DRP1 is required for force-induced mitochondrial fission. COS-7 cells were transfected with iLID-tdTomato-Omp25 and KIF5A-SspB. To suppress DRP1 activities, one group was treated with DRP1-targeted siRNA and another group with overexpression of DRP1(K38A). (E) Mitochondrial fission rates in the non-treated control group (NT), DRP1 siRNA treated group, and DRP1(K38A)-expressing group. N = 10 (NT), 11 (DRP1 siRNA), 13 (DRP1(K38A)) cells from 3 independent experiments. Statistical analysis using two-tailed paired Student’s t-test for the same condition, and two-tailed unpaired Student’s t-test for different conditions. (F) Fluorescence images of mitochondria in cells treated with DRP1 siRNA or DRP1(K38A) expression before and after blue light stimulation. Many mitochondria underwent force-induced tubulation (indicated by arrowheads), while few fission events occurred (indicated by yellow arrows). In (D) and (E), line, mean; bounds of box, 25th and 75th percentiles; whiskers, minimum to maximum value; circles, individual values. \*\*\**p* < 0.001, \*\*\*\**p* < 0.0001. Scale bars, (B-C) 2 μm, (F) 10 μm.

To further validate the involvement of DRP1, we checked whether and how DRP1 deficiency can influence force-induced fission. The activity of DRP1 in COS-7 cells was inhibited by the treatment of DRP1-targeted siRNA to knockdown DRP1, or the overexpression of a dominant negative DRP1 mutant, DRP1(K38A)^21^ which interacts with and consequently inhibits the GTPase activity of wildtype DRP1^22, 23^. In COS-7 cells expressing our optogenetic system, both methods generated many hyperfused mitochondria (Figure 4F), indicating the successful suppression of DRP1 functions. Next, we compared the fission rates before and during the 5 min intermittent blue light stimulation in the same cells. Blue light still triggered the drastic extension of thin mitochondrial tubules in cells treated with DRP1 knockdown or DRP1(K38A) overexpression (Figure 4F). However, few mitochondria divided following the force-induced mitochondrial elongation. Quantification of the mitochondrial fission rates showed that inhibition of DRP1, either by DRP1(K38A) overexpression (12.7×10^−3^ min^-1^ μm^-2^) or DRP1 knockdown (8.3 ×10^−3^ min^-1^ μm^-2^), significantly lowered the frequency of force-triggered fission compared to the control group (40.7 ×10^−3^ min^-1^ μm^-2^) (Figure 4E). To exclude the possibility that the long length of mitochondria is the key factor suppressing fission, we proved that long tubular mitochondria in COS-7 cells with functional DRP1 could still undergo force-triggered fission (Figure S4C). Therefore, our results show that DRP1 is required for mechanostimulation-induced mitochondrial fission.

### ER tubules cross over at force-induced mitochondrial fission sites

Endoplasmic reticulum (ER) has been identified as an important player in mitochondrial fission by utilizing ER tubules to wrap around and mediate the fission of mitochondria^24^. Here we examined whether ER tubules were present at mitochondrial fission sites upon forced-induced mitochondrial division (Figure 5A). COS-7 cells were transfected with iLID-tdTomato-Omp25 and KIF5A-SspB as well as GFP-Sec61β that marks the ER structures via the transmembrane domain of Sec61β, the β subunit of Sec61 complex localized on ER. Upon blue light-mediated mitochondrial elongation, the extending mitochondria tubule reached the network of ER. At the sites of fission either on the tubule (Figure 5B) or at the body-tubule junctions (Figure 5C), ER wrapped around the mitochondria right before the fission occurred. Moreover, we found that 81.25% of force-induced fission involved the ER wrapping around the fission sites (Figure 5D). Our results indicate that ER may participate in force-induced mitochondrial fission. More investigations are needed to uncover the mechanisms and exact roles of ER-mitochondria contact sites in the mechanoregulation of mitochondrial fission.

**Figure 5.**
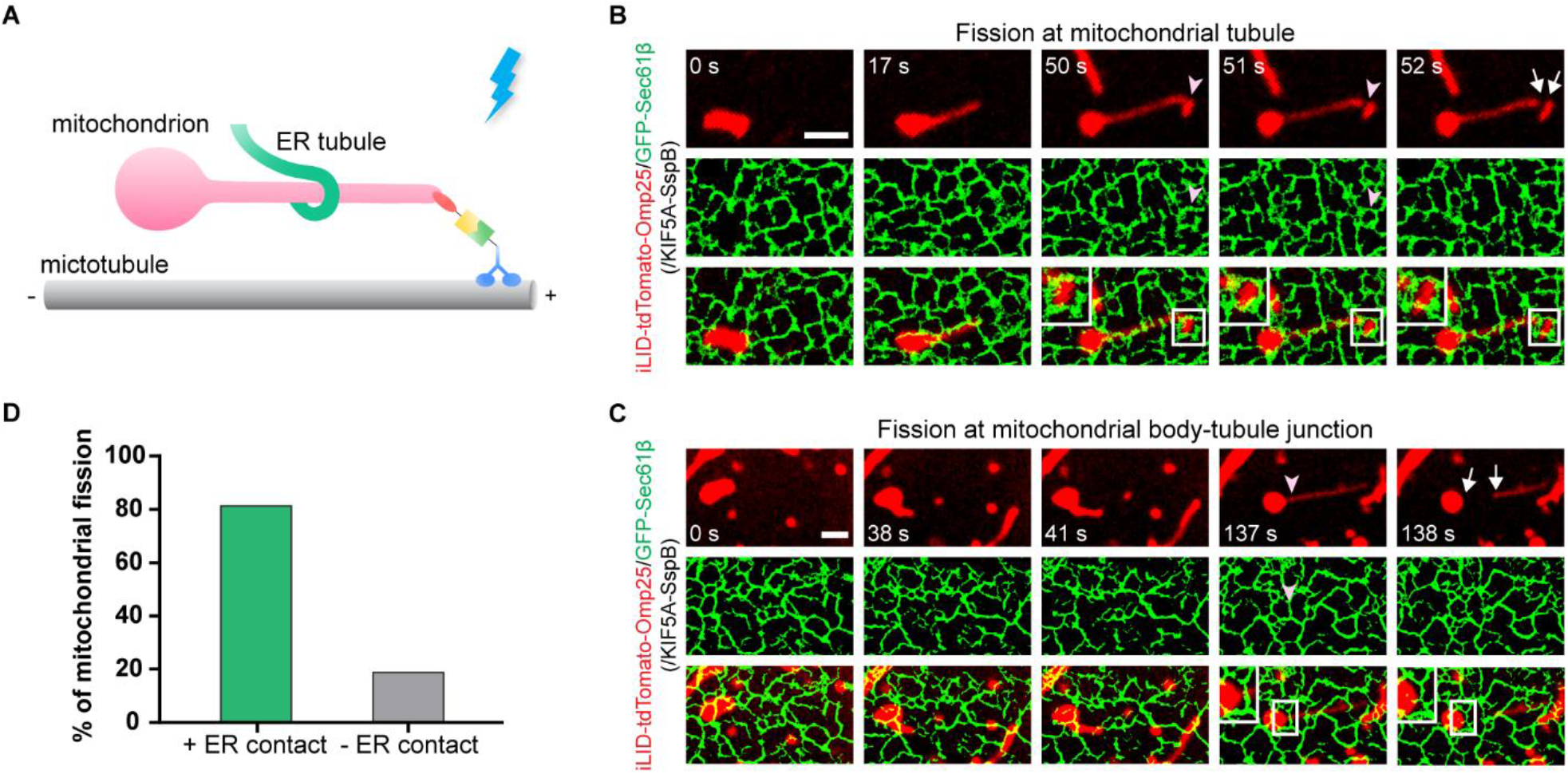
ER tubules cross over at force-induced mitochondrial fission sites. COS-7 cells were transfected with iLID-tdTomato-Omp25, KIF5A-SspB, and GFP-Sec61β. (A) Schematic illustration of ER tubules wrapping around the extending mitochondrial tubules to mediate force-induced fission. (B-C) Fluorescence images of ER and mitochondria. ER tubules wrapped the mitochondria at mitochondrial fission sites (indicated by pink arrowheads), either on the extending tubule (B) or at the mitochondrial body-tubule junctions (C). (D) Percentage of force-induced mitochondrial fission sites with or without ER contact at the force-induced fission sites. N = 32 fission events. Scale bars, 5 μm.

### Force-induced fission generates mitochondrial fragments without mtDNA which recruit Parkin proteins

We examined whether and how force-induced mitochondrial fission affects the distribution of mtDNA. mtDNA is the small and circular DNA residing in mitochondria that plays an essential role in maintaining mitochondrial functions^25^. To label mtDNA in living cells, TFAM, a mitochondrial transcription factor that packages mtDNA into nucleoids^26^, was fused with GFP. COS-7 cells were co-transfected with iLID-tdTomato-Omp25, KIF5A-SspB and TFAM-GFP. Intermittent blue light illumination for 5 min was delivered to cells to impose mechanostimulation towards mitochondria while the distribution of TFAM-GFP was monitored. During the forced-induced elongation and subsequent fission of mitochondria, in most cases, mtDNA stayed at its original intra-mitochondria location inside the mitochondria body (Figure 6A). As a result, mtDNA foci did not enter the extending tubules or the mitochondrial fragment disconnected from the body, irrespective of fission occurring on the tubule or the junction (Figure 6B, S5A). On a few occasions, mtDNA foci were redistributed into the extending tubules and entered the disconnected mitochondria fragment (Figure 6A, C, and S5B). Quantitative analysis shows that in 74.4% of force-induced mitochondrial fission, mtDNA did not enter the segregated mitochondrial tubules (Figure 6D). Moreover, we examined the fates of these disconnected mitochondrial fragments with or without mtDNA 300s after force-triggered fission (Figure 6E, S5C). For those with mtDNA, 39.4% of them could still undergo fission. Another 39.4% of them later fused with another mitochondrion containing mtDNA. On the contrary, for disconnected mitochondrial fragments without mtDNA, only 2% were divided afterward. 58.6% of them fused with other mitochondria. Intriguingly, mitochondrial fragments without mtDNA exhibited a strong preference to fuse with mitochondria with mtDNA (84.5%), which possibly serves to provide mtDNA complementation for further repair by fusion with mitochondria with normal mtDNA.

**Figure 6.**
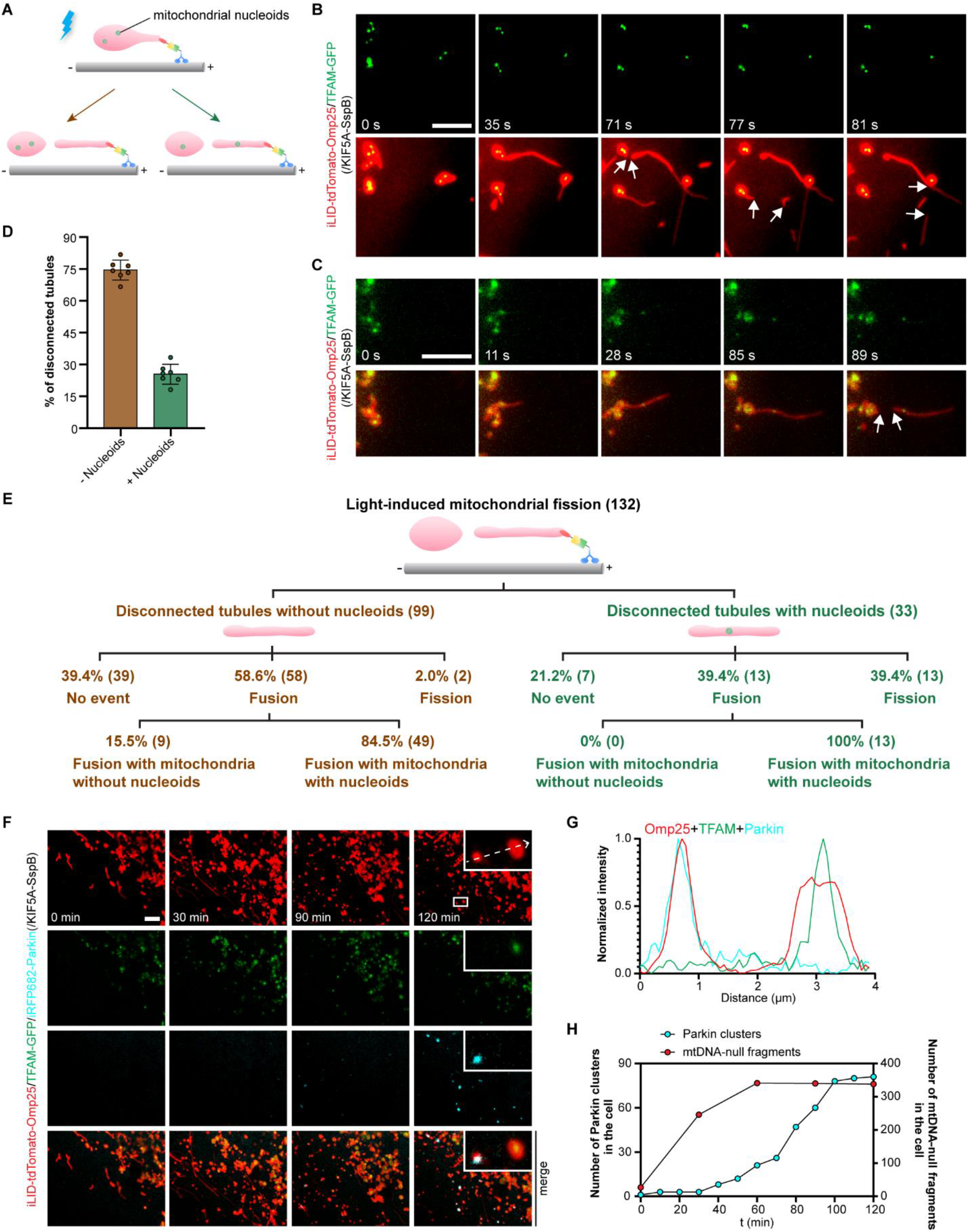
Force-induced fission generates mitochondrial fragments without mtDNA which recruit Parkin proteins. COS-7 cells were transfected with iLID-tdTomato-Omp25, TFAM-GFP and KIF5A-SspB (B-E) or along with iRFP682-Parkin (F-H). (A) Schematic illustration of the distribution of mitochondrial nucleoids after force-induced mitochondrial fission. (B-C) Fluorescence images of mitochondria and mtDNA. Upon force-induced fission (marked by white arrows), mtDNA stayed in the mitochondrial body (B) or got redistributed into the disconnected tubule (C). (D) The percentage of mitochondrial fission with disconnected tubules containing mtDNA. N = 227 fission events in total from 7 cells. Circles indicate values of individual cells. Data presents mean ± SD of ratio in individual cells. (E) Schema depicting the fates of the disconnected mitochondrial tubules which were tracked for 300 s after force-induced fission. (F) Fluorescence images of mitochondria, mtDNA and Parkin after intermittent blue light stimulation for 30 min and examined for 120 min. Tubules without mtDNA gradually recruited more Parkin proteins. (G) Plot profiles showing the fluorescence intensity of iLID-tdTomato-Omp25, TFAM-GFP and iRFP682-Parkin along the white dashed arrow. (H) The number of Parkin puncta and mtDNA-null fragments over time in the cell in (F). Scale bars, 5 μm.

Next, we asked what the fates are for mtDNA-null mitochondrial fragments which failed to fuse with another mitochondrion containing mtDNA. PINK/Parkin are core organizers in eliminating damaged mitochondria for mitochondrial quality control. Parkin amplifies a mitochondrial damage detection signal from PINK and recruits more Parkin to mitochondria. Here we tested whether Parkin can sense and mark mtDNA-null mitochondrial fragments generated by pulling force. Cells were transfected with iLID-tdTomato-Omp25, TFAM-GFP, KIF5A-SspB and iRFP682-Parkin and subjected to intermittent blue light illumination for 30 min. Around 90 min after the start of blue light exposure, discernable Parkin clusters formed exclusively on mtDNA-null mitochondria and increased over time (Figure 6F, whole cell images in Figure S6). The examination of the fluorescence intensity along the white dashed arrow confirmed the colocalization of Parkin clusters and mtDNA-null fragments (Figure 6G). We quantified the number of Parkin clusters at different time points and confirmed its gradual increase along with the formation of mtDNA-null fragments over time (Figure 6H). Our results link the force-induced mitochondrial fragments containing no mtDNA with mitophagy and mitochondrial quality control. Mechanisms underlying the preference of mtDNA remaining inside the mitochondria body upon mechanostimulation and the role of Parkin in the fate of force-induced mtDNA-null mitochondrial fragments await more investigation.

## Discussion

In this study, we used an optogenetic mitochondria-specific intracellular mechanostimulator to interrogate the connections between mechanical forces and mitochondria dynamics. We found that while imposing pulling force on mitochondria can induce the protrusion of membrane tubules from the mitochondrial body and the subsequent fission, sustained force exertion can promote fission more effectively than transient application. The force-induced fission can occur either at the body-tubule junctions or at tubules, which also leads to the asymmetrical segregation of mtDNA among the body and the disconnected tubule fragment. DRP1 and ER tubule wrapping were involved in the force-triggered mitochondrial fission. Moreover, we revealed that the daughter mitochondrial fragments containing mtDNA would recruit Parkin proteins. Our results not only directly prove the mechanosensitive and mechanoresponsive nature of mitochondria but also reveal the direct mechanoregulation of fission, placing mitochondria as a mechanosensor on the complex map of cellular mechanosensing and mechanotransduction.

We found that mitochondrial division upon force application always occurred at the protruding tubules that possess distinct morphological features compared with the body. A previous report has shown that higher mitochondrial membrane tension promotes mitochondrial division^17^. In our experiments, the thin mitochondrial tubule pulled from the mitochondrial body has higher local membrane tension due to its smaller diameter, which may account for the preference for fission at the tubules. The protrusion of long mitochondrial tubules also increases the likelihood of reaching adjacent ER structures and generating more ER-mitochondria contact sites, which may also promote mitochondrial fission. In fact, the tubulation of mitochondria naturally occurs in cells by intracellular mechanical stimulation (e.g., the pulling forces provided by molecular motors or associated organelles). Here our work indicates the direct contribution of mitochondrial elongation to driving mitochondrial fission, in addition to its previously reported roles, including promoting the formation of the mitochondrial network^4^ and regulating the transportation of mtDNA^27^. Intriguingly, we also uncovered that force-induced deformation induces the ring closure of donut-shaped mitochondria. Taken together, our results highlight the role of mechanical forces in sculpting mitochondrial shapes.

Our results also consolidate the mechanosensitivity and mechanoresponsiveness of mitochondria. While tremendous efforts have been dedicated to uncovering the participation of the plasma membrane, cytoskeleton, and various intracellular signaling pathways in cellular mechanotransduction, increasing interest has been drawn to various organelles. For example, a recent seminal report brought to light the mechanosensitivity of Golgi and showed that Golgi could respond to mechanical cues by modulating lipid metabolism^28^. Moreover, two decades of endeavors have identified the cell nucleus as a critical mechanosensor^29^. Here our results prove that mitochondria can directly sense and respond to mechanical forces, providing direct proof for the role of mitochondria as an important player in the complex network of cell mechanosensing and mechanotransduction. As mitochondria execute diverse functions, more investigations are needed to elucidate how other mitochondrial functions can be directly modulated by mechanostimulation. Therefore, it would be very interesting to study how mechanical stimuli influence cell functions via the mechanoregulation of mitochondrial dynamics in various physiological and pathological settings.

Moreover, we showed that sustained mechanostimulation induced a large number of mitochondrial fission events and triggered Parkin-dependent mitochondrial quality control. In cells, excessive fission results in the imbalance of mitochondrial homeostasis and causes mitochondrial fragmentation. Fragmentation of the mitochondria leads to mitochondrial bioenergetics deficits and has been implicated in many pathological states. Indeed, intensive or acute mechanical stress has been shown to induce mitochondrial fragmentation in vivo in previous reports^11,12^. Therefore, our results raised the possibility that mechanical over-stimulation, by either external or intracellular forces (e.g., abnormal motor activities or hyper-active organelle hitchhiking), poses severe stress to cells via the pathway of force-induced mitochondria hyper-fission.

In summary, utilizing optogenetic stimulation of intracellular mitochondria, we found that mechanical force exerted on mitochondria in cells can induce mitochondrial fission which involves DPR1 and ER tubule wrapping. The force-induced fission generates daughter mitochondrial fragments containing no mtDNA that accumulate Parkin. Our results provide new insights into mitochondrial dynamics and mechanobiology, paving the road to future investigation of mechanoregulation of mitochondrial functions in health and disease.

## Methods

### Cell cultures

COS-7 (ATCC® CRL-1651™) cells were grown in DMEM medium (Thermo Fisher Scientific) supplemented with 10% FBS (fetal bovine serum, Clontech) and 1% P/S (Penicillin-Streptomycin, Thermo Fisher Scientific). U2OS (ATCC® HTB-96™) cells were cultured in McCoy’s 5A medium (Thermo Fisher Scientific) supplemented with 10% FBS (fetal bovine serum, Clontech) and 1% P/S (Penicillin-Streptomycin, Thermo Fisher Scientific). All cultures were maintained in a humidified environment at 37°C with 5% CO_2_.

### Plasmid construction

The SspB variant named micro was used in our study. KIF5A-GFP-SspB, KIF1A-GFP-SspB, ePDZ-mCh-Miro1, KIF5A-GFP-LOVpep, CRY2-mCh-Miro1, and KIF5A-GFP-CIBN have been described in our previous work^14, 30^. KIF5A-SspB sequence was inserted into the pEGFPN1 backbone to make KIF5A-SspB. mCh-DRP1 was obtained from Addgene (Addgene #49152), and GFP-DRP1 was then constructed by replacing mCherry with GFP using In-Fusion (Clonetech). mCh-Omp25 (a.a 111-145), DRP1K38A, TFAM-dsRed, mCh-Parkin were kindly provided by Prof. LIU Xingguo from Guangzhou Institutes of Biomedicine and Health, Chinese Academy of Sciences. DRP1(K38A) and TFAM sequences were inserted into mammalian expression pEGFPN1 vector to construct GFP-DRP1(K38A) by In-Fusion (Clonetech) and TFAM-GFP by ligation. By In-Fusion (Clonetech), iLID-tdTomato-Omp25 was obtained by inserting iLID and Omp25 into the backbone containing tdTomato. Using iLID-tdTomato-Omp25 as a backbone, iLID-tdTomato was replaced by tdTomato to make tdTomato-Omp25 using ligation. Using In-Fusion (Clonetech), CRY2-tdTomato-Miro1 was made by inserting CRY2, tdTomato and Miro1 into pEGFPN1 backbone, and iLID and tdTomato-Miro1 were fused into pEGFPN1 backbone to obtain iLID-tdTomato-Miro1. GFP-Sec61β (a.a 1-96) was cloned by inserting Sec61β into pEGFPN1 vector using ligation. Parkin gene was inserted into iRFP682-Smad2 (Addgene #118943) to construct iRFP682-Parkin. Restriction enzymes from Invitrogen™ were used for all the linearization of vectors and CloneAmp™ HiFi PCR Premix (Takara) was used for all the PCR reactions. Details for plasmid constructions are listed in Table S1.

### Plasmid transfection and siRNA transfection

Cells were plated on Poly-D-Lysine coated 35mm confocal dishes with a hole size of 13Ø 24h. Cells were then transfected by Lipofectamine™ 3000 Reagent (Thermo Fisher Scientific) according to the manufacturer’s protocol and recovered overnight before imaging. DRP1 targeted small interfering RNAs (sense strand: 5′-UCCGUGAUGAGUAUGCUUUdT dT-3′) as previously described^31^ were purchased from Sangon Biotech (Sangon, Shanghai, China) for the knockdown of DRP1. Transfection of siRNA was performed using Lipofectamine™ RNAiMax Reagent (Thermo Fisher Scientific), and cells were analyzed 48∼72h after siRNA transfection.

### Live cell imaging

The fluorescence imaging of mitochondria, mitochondrial nucleoids, and DRP1 puncta was performed on an epifluorescence microscope (Leica DMi8S, Thunder Imager) equipped with an on-stage CO_2_ incubator using a 100× oil immersion objective. Images were captured with the following excitation/emission filter settings: 470nm/510nm for GFP, 575nm/590nm for mCh and tdTomato, 640nm/700nm for iRFP682. Blue light illumination at 480 mW/cm^2^ was used to excite both optogenetic pairs and GFP. Pulsed blue light with 200ms exposure duration was given to the cells at 1-s intervals when sustained blue light stimulation was needed. The spatial control of blue light in ROI was realized by the infinity scanner on Leica DMi8S. Two-color live imaging of GFP-Sec61β and iLID-tdTomato-Omp25 was performed on a spinning disk confocal microscope (Leica DMi3000B) with 488nm and 561nm lasers using a 100× oil immersion objective. For this experiment, 561nm laser (200 ms exposure, laser power 10%-20%) was utilized to excite and image tdTomato, and 488nm laser (200 ms exposure, laser power 10%-20%) was used for blue light stimulation and GFP visualization. The images of ER were later processed to increase the contrast and resolution using a previously reported algorithm^32^.

### Measurement of mitochondrial fission rate

In live cells, the division of one single mitochondrion into two daughter mitochondria was counted as one mitochondrial fission event. Taking the previously published method^33^ as a reference, the fission rate of mitochondria in our study was defined as the number of fission events per square micrometer area of mitochondria. To measure the area of mitochondria, images were segmented by the trainable Weka segmentation (Fiji plugin) and then binarized. The mitochondrial area in binarized images was measured by using ImageJ Thresholding. The data was plotted and analyzed by Graphpad Prism.

### Characterization of mitochondrial fission sites

Image analysis of mitochondrial fission sites was performed with ImageJ/Fiji. For differentiating different fission sites on mitochondria, mitochondrial tubule was recognized as the thin, tubular part which was stretched out from mitochondria, and the junction was indicated as the small region on the tubule close to the spherical mitochondria body (<300nm).

### Measurement of GFP-DRP1 intensity

GFP-DRP1 signal was quantified in the raw fluorescence images using ImageJ/Fiji. As described previously^33^, the intensity of GFP-DRP1 was measured by drawing a ∼400nm circle at the fission site on mitochondria, and the GFP-DRP1 intensity of the same sized circle at non-fission sites on the same mitochondrion was measured. The datasets were scaled by dividing by the mean value of GFP-DRP1 intensity of non-fission sites corresponding to fission at the junction. The graphs of GFP-DRP1 intensity were created by Graphpad Prism.

### Line profile plots

All the line profile plots were created by measuring the raw fluorescence images. Line profiles were analyzed by ImageJ/Fiji, and graphs were made by Graphpad Prism. The intensity was normalized by the formula (F-F_min_)/(F_max_-F_min_), where F refers to fluorescence intensity, and F_min_ and F_max_ are the minimum and maximum fluorescence intensity respectively along the line.

### Statistics

All the statistical analyses were performed using Graphpad Prism. Normality test was conducted for each dataset using D’Agostino-Pearson normality test (p>0.05). For datasets considered as normally distributed, a two-sided t-test was selected for comparing two groups. Datasets which failed the normality test were analyzed by a nonparametric test (Wilcoxon Matched-Pairs signed ranks test). The p values were indicated in the graphs: ns p>0.05, \**p* < 0.05, \*\**p* < 0.01, \*\*\**p* < 0.001, \*\*\*\**p* < 0.0001.

## Acknowledgments

We thank Prof. Liwen Jiang for equipment support and helpful discussion.

## Author contributions

L.D., X.L. conceived the project and designed experiments. X.L., L.X., Y.S., and X.L. performed the experiments. All the authors discussed and analyzed the data. L.D., X.L., Y.S., L.X. designed figures. L.D. and X.L. wrote the manuscript with input from all authors.

