## Supplementary materials for "Mechanical force induces DRP1-dependent asymmetrical mitochondrial fission for quality control"

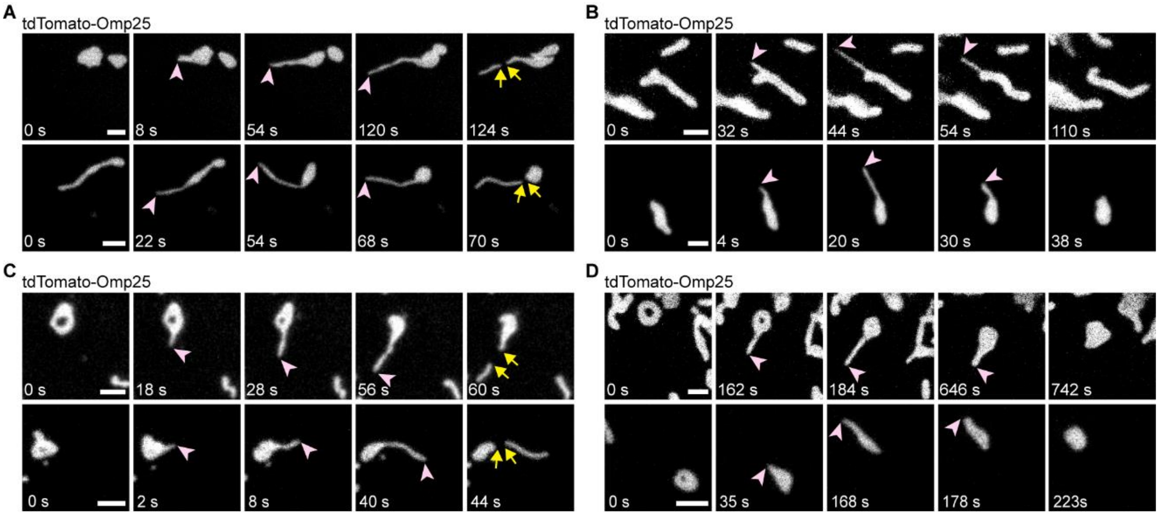


**Figure S1. The naturally occurring mitochondrial tubulation alters the shape of mitochondria and induces fission.** COS-7 cells were transfected with tdTomato-Omp25. (A) After mitochondrial tubulation, mitochondrial fission occurred either at the body-tubule junction or on the tubule. (B) Upon mitochondrial tubulation, some tubules were withdrawn to the mitochondrial body. (C) The mitochondrial tubulation caused the ring closure of the donut-shaped mitochondria, followed by fission at different sites. (D) Upon tubulation, the donut-shaped underwent ring closure and the extending tubule was retrieved to the body. Scale bars, 2 μm.

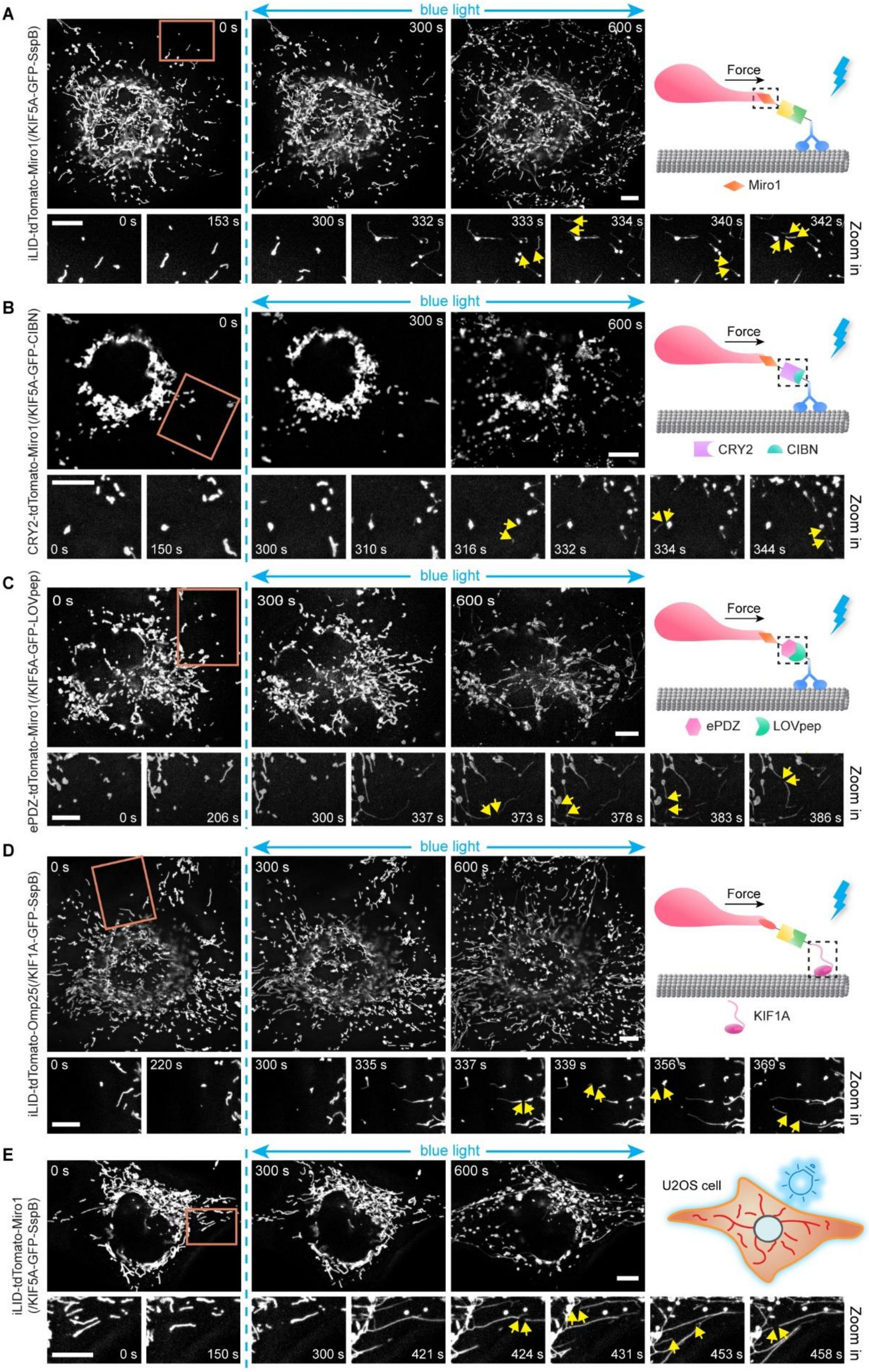


**Figure S2. Force-induced mitochondrial fission can be achieved with different mitochondrial targeting sequences, optical hetero-dimers, motor proteins, and in other types of cells.** COS-7 cells were subjected to 5 min intermittent blue light stimulation and the mitochondria dynamics were monitored 5 min before and after the start of blue light illumination. (A) Fluorescence images of mitochondria in COS-7 cells expressing iLID-tdTomato-Miro1 and KIF5A-GFP-SspB, where Miro1 is another mitochondrial outer membrane targeting sequence. (B-C) Fluorescence images of mitochondria in COS-7 cells expressing CRY2-tdTomato-miro1+KIF5A-GFP-CIBN (B), or ePDZ-tdTomato-miro1+KIF5A-GFP-LOVpep (C), where CRY2/CIBN and ePDZ/LOVpep are another two pairs of blue light-gated optical dimerizers. (D) Fluorescence images of mitochondria in COS-7 cells expressing iLID-tdTomato-Omp25+ KIF1A-GFP-SspB, where KIF1A encodes another type of kinesin motor, kinesin 3. (E) Fluorescence images of mitochondria in U2OS cells expressing iLID-tdTomato-Miro1 and KIF5A-GFP-SspB. Scale bars, 10 μm.

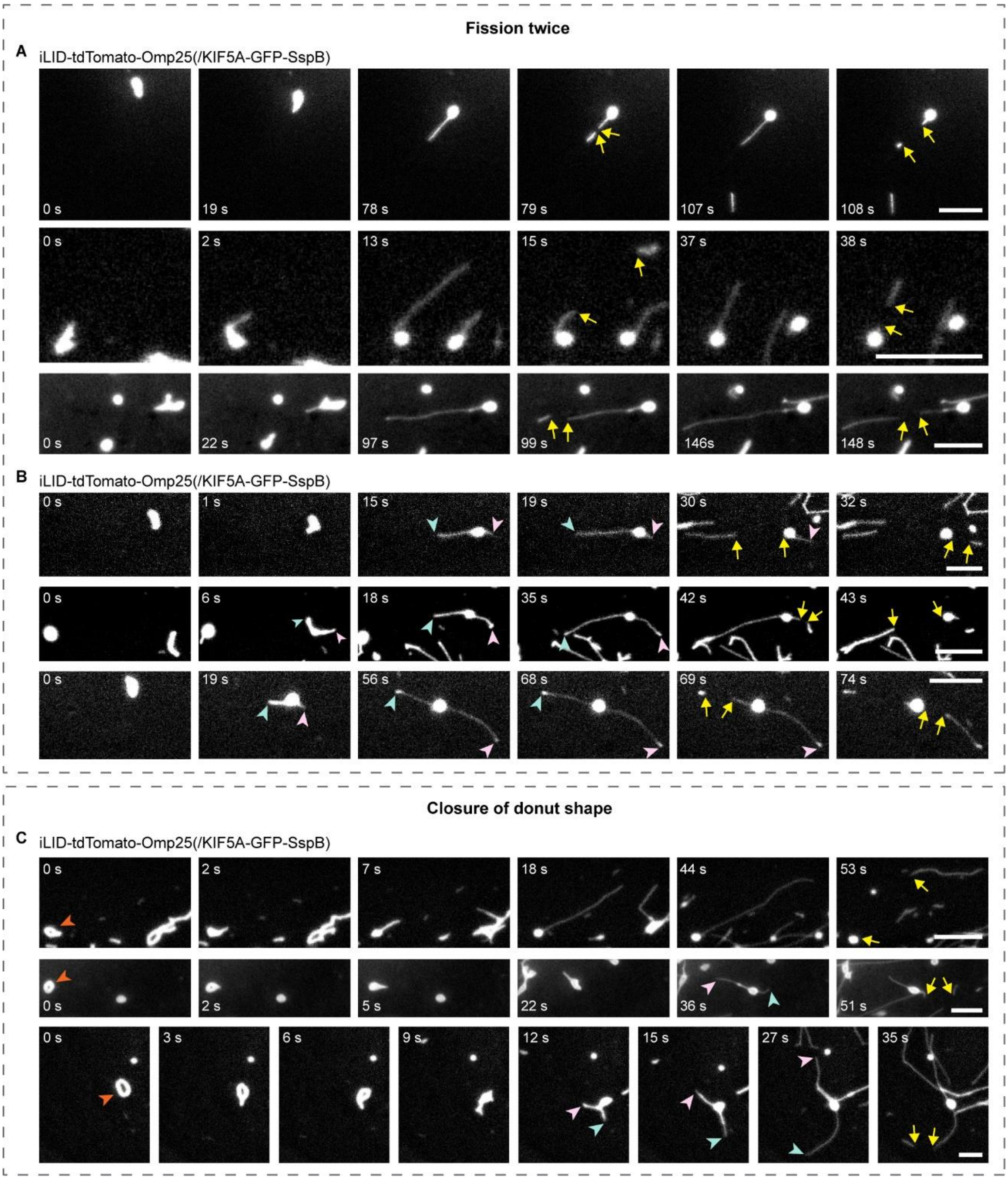


**Figure S3. Additional examples for different phenomena during force-induced mitochondrial fission.** COS-7 cells were transfected with iLID-tdTomato-Omp25 and KIF5A-GFP-SspB. (A-B) Fluorescence images of mitochondria undergoing force-induced fission twice. (A) Fission occurred twice at the same extending tubule. (B) Fission occurred at either of the two tubules protruding from the same mitochondria. (C) Fluorescence images of donut-shaped mitochondria (indicated by orange arrowheads) undergoing ring closure before force-induced fission. Scale bars, 5 μm.

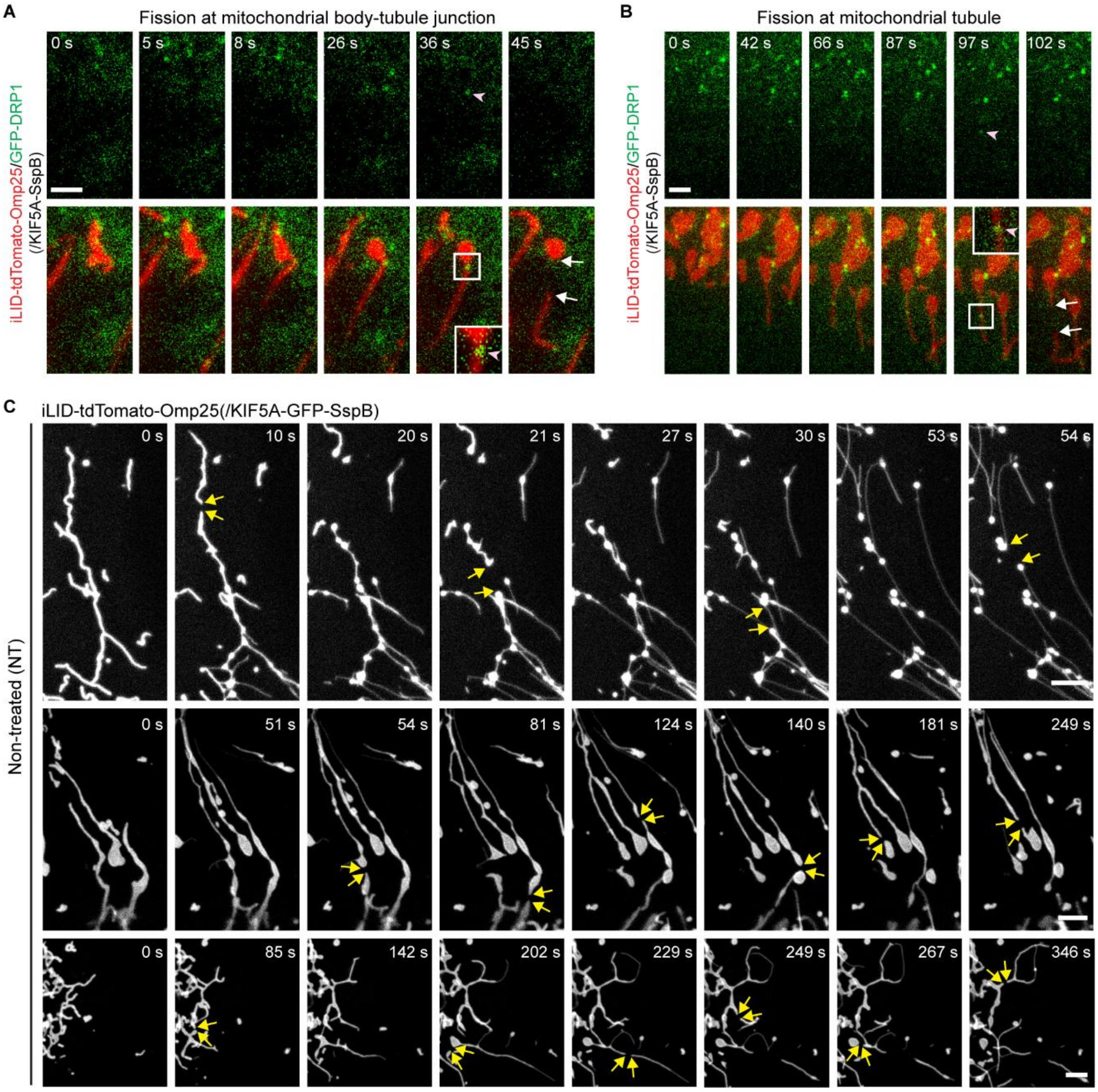


**Figure S4. DRP1 participates in mechanical force-induced mitochondrial fission.** (A-B) Additional examples for DRP1 recruited on either the junction (A) or the tubule (B) during force-induced mitochondrial fission. COS-7 cells were transfected with iLID-tdTomato-Omp25, GFP-DRP1 and KIF5A-SspB. (C) Long tubular mitochondria in COS-7 cells with functional DRP1 could still undergo force-triggered fission. COS-7 cells were transfected with iLID-tdTomato-Omp25, and KIF5A-GFP-SspB without any treatment to suppress DRP1 activities. Scale bars, (A-B) 2 µm, (C) 5 µm.

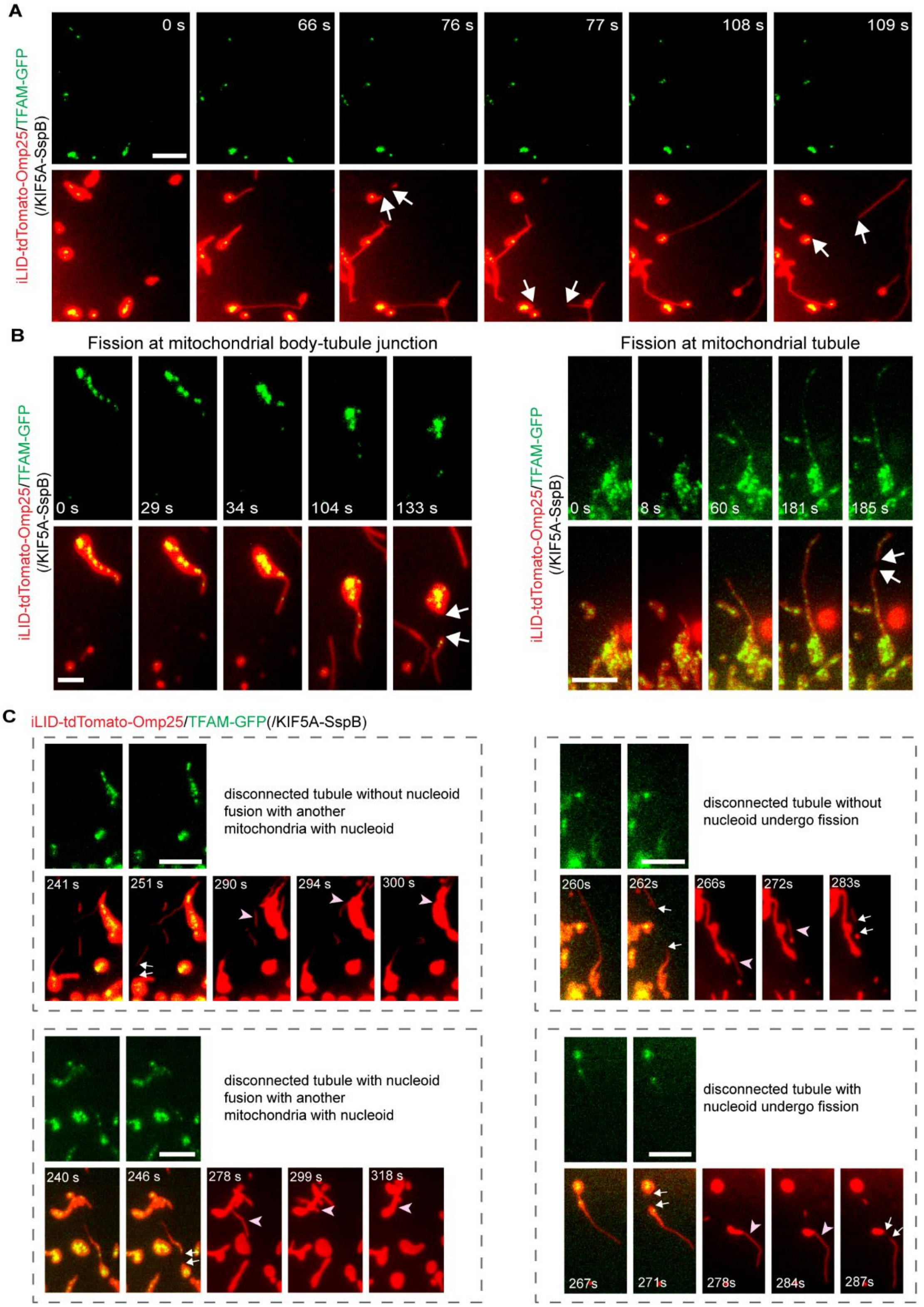


**Figure S5. Asymmetrical distribution of mtDNA during force-induced fission.** COS-7 cells were transfected with iLID-tdTomato-Omp25, TFAM-GFP and KIF5A-SspB. (A-B) Additional examples of mtDNA distribution during force-induced fission (indicated by white arrows). (A) During the force-induced fission at the body-tubule junction, mtDNA was not distributed in the disconnected tubule. (B) During the force-induced fission at the body-tubule junction or on the tubule, mtDNA was distributed into the disconnected tubule. (C) Examples of disconnected tubules with different fates after force-induced fission. In the example showing disconnected tubule without nucleoid undergo fission, field of view is moved up in between the image at time point 262s and the image at 266s to track the disconnected tubule. Scale bars, 5 μm.

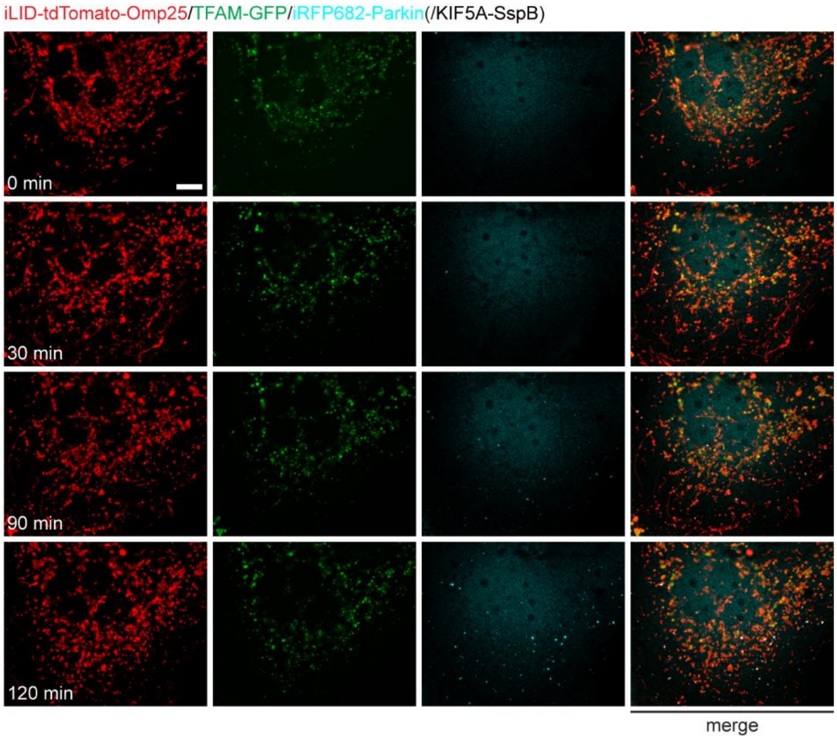


**Figure S6. Force induces the formation of mtDNA-null mitochondrial fragments which recruit Parkin proteins.** COS-7 cells were transfected with iLID-tdTomato-Omp25, TFAM-GFP, KIF5A-SspB and iRFP682-Parkin. Fluorescence images of the whole cell from Figure 6F. Scale bars, 10 μm.

**Tables**

**Table S1. Plasmids construction.**

| Plasmids | Method | Template | Insertion sites | Forward Primer | Reverse Primer |
|---|---|---|---|---|---|
| GFP-DRP1 | In-Fusion | mCh-DRP1 | BshTI and XhoI | cgctagcgctaccggtgccaccatggtgagcaag | gcctccatcactcgagacttatacagctcgtccatgcc |
| CRY2-tdTomato-Miro1 | In-Fusion | pEGFPN1 | Bsp1407I and MunI | atggacgagctgtacaagcc | ttaacaacaacaattgcattcattttatgtttcagg |
| iLID-tdTomato-Miro1 | In-Fusion | pEGFPN1 | BshTI and HindIII | tggaccggagaccggtatggtgagcaagggcgaggag | gcagaattcgaagctagttatctagatccggtggatccc |
| GFP-DRP1(K38A) | In-Fusion | GFP-DRP1 | XhoI and BamHI | ggactcagatctcgaggcatggaggcgctaattcctgt | tagatccggtggatcctcaccaaagatgagtctcccgga |
| iLID-tdTomato-Omp25 | In-Fusion | iLID-tdTomato-Sec61β | Bsp1407I and NotI | atggacgagctgtacaagtccg | tctagagtcgcggccgcccccccctttttctggagactaaataaaatc |
| TFAM-GFP | Ligation | pEGFPN1 | NheI and BshTI | - | - |
| iRFP682-Parkin | Ligation | iRFP682-Smad2 | Bsp1407I and MunI | - | - |
| tdTomato-Omp25 | Ligation | iLID-tdTomato-Omp25 | NheI and Bsp1407I | - | - |
| GFP-Sec61β | Ligation | pEGFPN1 | Bsp1407I and NotI | - | - |
